# Developing snakebite risk model using venomous snake habitat suitability as an indicating factor: An application of species distribution models in public health research

**DOI:** 10.1101/2020.04.06.027342

**Authors:** Masoud Yousefi, Anooshe Kafash, Ali Khani, Nima Nabati

## Abstract

Snakebite envenoming is an important public health problem in Iran, despite its risk not being quantified. This study aims to use venomous snakes’ habitat suitability as an indicator of snakebite risk, to identify high-priority areas for snakebite management across the country. Thus, an ensemble approach using five distribution modeling methods: Generalized Boosted Models, Generalized Additive Models, Maximum Entropy Modeling Generalized Linear Models, and Random Forest was applied to produce a spatial snakebite risk model for Iran. To achieve this, four venomous snakes’ habitat suitability (*Macrovipera lebetina, Echis carinatus, Pseudocerastes persicus* and *Naja oxiana*) were modeled and then multiplied. These medically important snakes are responsible for the most snakebite incidents in Iran. Multiplying habitat suitability models of the four snakes showed that the northeast of Iran (west of Khorasan-e-Razavi province) has the highest snakebite risk in the country. In addition, villages that were at risk of envenoming from the four snakes were identified. Results revealed that 51,112 villages are at risk of envenoming from *M. lebetina*, 30,339 from *E. carinatus*, 51,657 from *P. persicus* and 12,124 from *N. oxiana*. This paper demonstrates application of species distribution modeling in public health research and identified potential snakebite risk areas in Iran by using venomous snakes’ habitat suitability models as an indicating factor. Results of this study can be used in snakebite and human–snake conflict management in Iran. We recommend increasing public awareness of snakebite envenoming and education of local people in areas which identified with the highest snakebite risk.

## Introduction

Snakebite envenoming is known as an important public health problem and the cause of medical emergencies around the globe [1-14]. On Earth, between 421,000 and 1.2 million people are envenomed by venomous snakes annually and around 125,000 deaths per year are attributable to snakebite envenoming [1, 6, 14, 15]. Snakebite envenoming is mostly described as a neglected public health issue in the tropics, [2, 6, 16, 17] however, it is also an important challenge for public health in temperate areas like Iran [18].

Iran is home to 81 snake species [19, 14] of which 25 are venomous (Nine species are sea snakes and 16 species are terrestrial snakes). *Macrovipera lebetina, Echis carinatus, Pseudocerastes persicus* and *Naja oxiana* are widespread in Iran and are responsible for the most snakebite incidents in the country [18, 21]. A study reported 53,787 cases of snake bites between 2002 and 2011 in Iran [18]. Despite considerable research into the phylogeny, taxonomy, morphology and ecology of venomous snakes in Iran [22, 31] snakebite envenoming has received less attention [18, 32]. In fact, snakebite is an important uninvestigated public health problem and conservation challenge in Iran [33-35]. Thus, more effort should be made to identify areas with high snakebite risk and reduce envenoming risk from snakes.

Species distribution models (SDMs) have found an important application in biodiversity research [36-38]. They are employed in studying habitat suitability [39-41], identifying environmental drivers of species distribution [42-45] and predicting impacts of climate change on biodiversity [46-51]. SDMs are successfully used to identify suitable habitats of species even in areas with no distribution records [52-54]. Thus, these models can be used to identify suitable habitats of venomous snakes as proxies of snakebite risk [12, 55-57] in data poor regions like Iran.

The main goal of this paper was to apply SDMs and produce a spatial risk model for snakebite in Iran. To do this, firstly, five distribution modeling methods [38]: Generalized Boosted Models, Generalized Additive Models, Maximum Entropy Modeling Generalized Linear Models, and Random Forest and distribution data of *M. lebetina, E. carinatus, P. persicus* and *N. oxiana* were to produce their habitat suitability models. Secondly, the five habitat suitability models [38-58] of each species were combined by ensemble approach and finally the four species ensemble models were multiplied to identify potential snakebite risk. We also in addition, number of villages that are at risk of envenoming by these four snakes determined in Iran.

## Materials and methods

### Occurrence data

Distribution records of the *M. lebetina, E. carinatus, P. persicus* and *N. oxiana* (Fig 1) were obtained from published books and papers [31, 59-66], and online databases (GBIF and VertNet). These four snakes were selected because they are responsible for the most snakebite incidents in Iran [18] and have the widest distribution range across the country [19-20]. By combining presence records from the two sources 89 distribution records were obtained for *M. lebetina*, 68 records for *E. carinatus*, 54 records for *P. persicus* and 37 records for *N. oxiana* (Fig 2).

**Fig 1.**
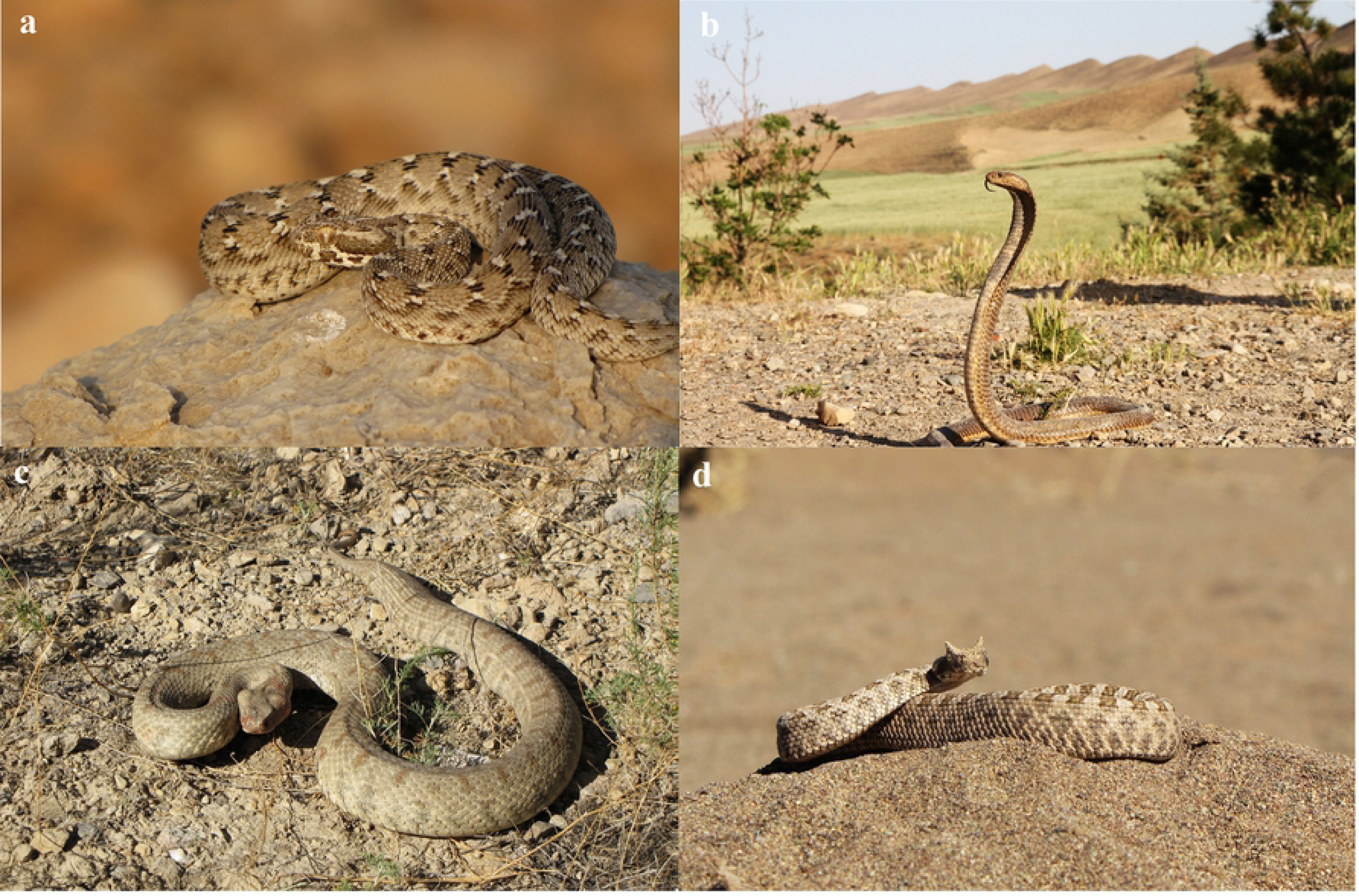
Photos of the four medically important venomous snakes modeled in this research to map snakebite in Iran. Saw-scaled Viper, *Echis carinatus* (a), Central Asian Cobra, *Naja oxiana* (b), Levantine Viper, *Macrovipera lebetina* (c), and Persian Horned Viper, *Pseudocerastes persicus* (d). Photos by Masoud Yousefi.

**Fig 2.**
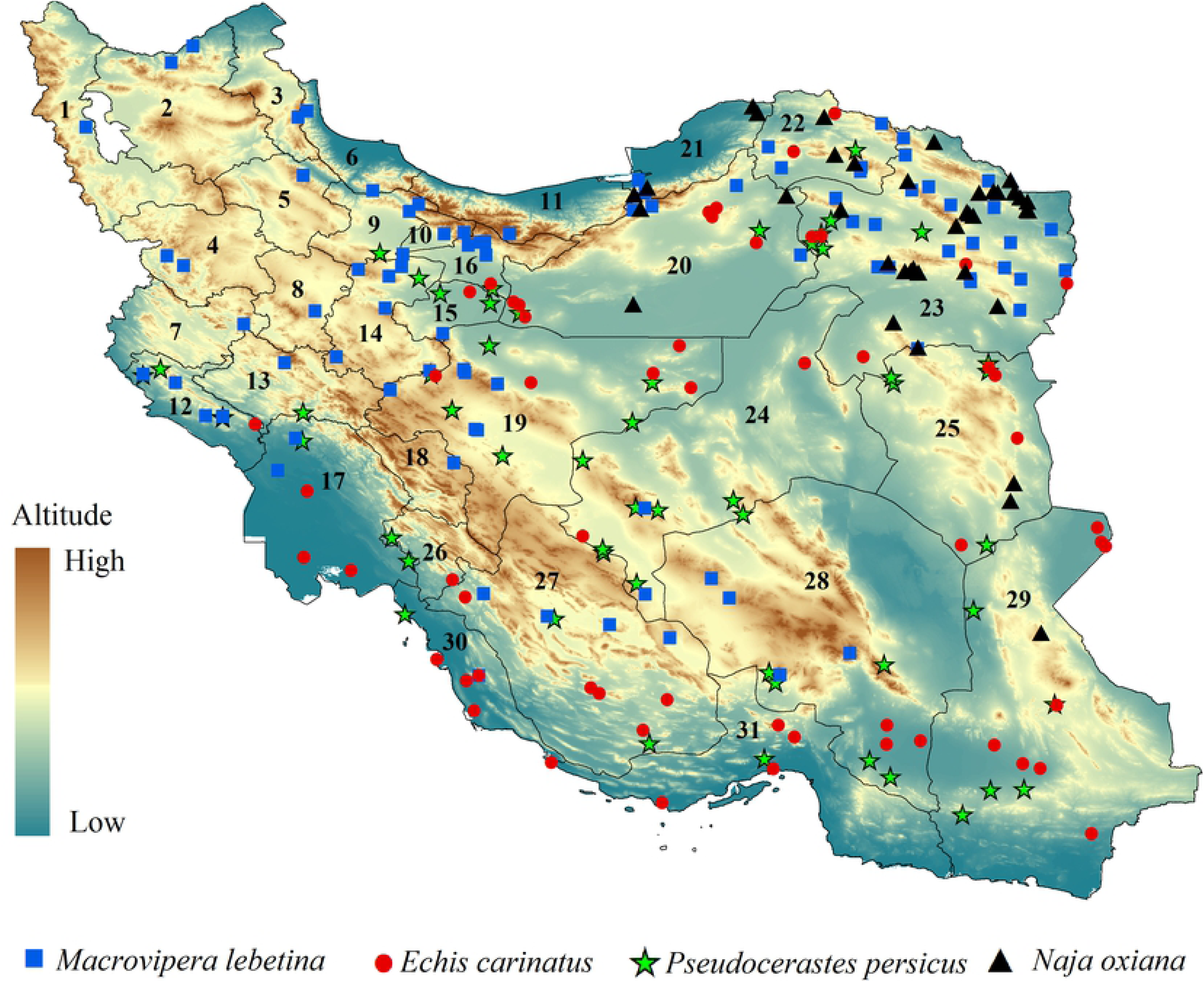
Distribution records of *Macrovipera lebetina, Echis carinatus, Naja oxiana*, and *Pseudocerastes persicus* in a topographic overview of Iran. Provinces numbers should read as follows; (1) West Azerbaijan, (2) East Azerbaijan, (3) Ardabil, (4) Kurdistan, (5) Zanjan, (6) Gilan, (7) Kermanshah, (8) Hamedan, (9) Qazvin, (10) Alborz, (11) Mazandaran, (12) Ilam, (13) Lorestan, (14) Markazi, (15) Qom, (16) Tehran, (17) Khuzestan, (18) Chahar Mahaal and Bakhtiari, (19) Isfahan, (20) Semnan, (21) Golestan, (22) Khorasan-e-Shomali (Northern Khorasan), (23) Khorasan-e-Razavi, (24) Yazd, (25) Khorasan-e-Jonobi (Southern Khorasan), (26) Kohgiluyeh and Boyer-Ahmad, (27) Fars, (28) Kerman, (29) Sistan and Baluchestan, (30) Bushehr, (31) Hormozgan.

### Environmental data

Seven uncorrelated environmental variables (Table 1) related to climate, topography, vegetation, and human footprint were used to develop the four snakes’ habitat suitably models [12, 17, 28, 55, 67-70]. Climatic variables were downloaded from the WorldClim database at 30-seconds spatial resolution [71]. Normalized Difference Vegetation Index (NDVI) was considered as an indicator of resource availability [72]. Snakebite risk is associated with human population density and activities [55], thus, human footprint index was included in models [69]. Human footprint index was produced by combining data on the extent of built environments, population density, electric infrastructure, crop lands, pasture lands, roads, railways, and navigable waterways [70]. Topographic heterogeneity was used as topography variable by measuring the standard deviation of elevation values in area grid cells of 1km from a 90 m resolution in the Raster package [73] in the R environment v.3.4.3 [74]. Elevation layer was obtained from the Shuttle Radar Topography Mission (SRTM) elevation model [75].

**Table 1.** List of Environmental variables. Name, source and the variance inflation factor (VIF) for the seven environmental variables which were used for developing habitat suitability of venomous snakes in Iran.

| Variable | Description (abbreviation) | References | VIF |
| --- | --- | --- | --- |
| Climate | Temperature seasonality (bio4) | [71] | 1.438 |
|  | Annual precipitation (bio12) | [71] | 2.871 |
|  | Precipitation seasonality (bio15) | [71] | 2.389 |
|  | Precipitation of driest quarter (bio17) | [71] | 3.049 |
| Topography | Topographic heterogeneity (SD) | [75] | 1.315 |
| Vegetation | Normalized Difference Vegetation Index (NDVI) | [72] | 3.33 |
| Anthropogenic | Human Footprint (HFP) | [69-70] | 1.466 |

### Snakebite risk modeling

An ensemble approach [38-58] was applied to model habitat suitability of the *M. lebetina, E. carinatus, P. persicus* and *N. oxiana*, using five methods: Generalized Boosted Models (GBM; [76]), Generalized Additive Models (GAM; [77]), Maximum Entropy modeling (Maxent; [78]), Generalized Linear Models (GLM; [79]), Random Forest (RF; [80]). Since these methods need background data points we generated a randomly drawn sample of 10,000 background points (e.g., pseudo-absence points) from the extent of the study area using the PresenceAbsence package [81]. Then areas associated with high snakebite risk in Iran were identified by multiplying habitat suitability models of the four species.

To quantify snakebite risk in Iran in more detail number of villages that are at risk of envenoming from the four snakes determined and area of snakebite risk calculated in the Raster package [73]. For this continuous habitat suitability models were converted to suitable/unsuitable maps using maximum test sensitivity with a specificity threshold [82]), and then overlaying 185,000 (population in these villages range from less than 50 to 5000 individuals) villages with each snake model.

### Model performance

Assessing model performance is an important step in distribution modeling studies [37]. Several metrics were introduced for estimating the predictive powers of models [37, 78, 83, 84]. Three metrics were considered to assess the produced habitat suitability models’ performances. The true skills statistic (TSS), area under the receiver operating characteristic curve (AUC), and the Boyce index [37, 83, 85].

Analyses were carried out using the packages GISTools (https://rdrr.io/cran/GISTools/), dismo (https://rdrr.io/cran/dismo/), biomod2 (https://cran.rproject.org/web/packages/biomod2/index.html), maptools (http://r-forge.r-project.org/projects/maptools/), SDMTools (htt ps://cran.r-project.org/web/packages/SDMTools/index.html), and ecospat (https://cran.rproject.org/web/packages/ecospat/ecospat.pdf), in the R environment (v.3.4.3).

## Results

### Model performance

All models developed in this study performed well based on the three model performance evaluation metrics, AUC, TSS and Boyce index (See figs 3-6).

**Fig 3.**
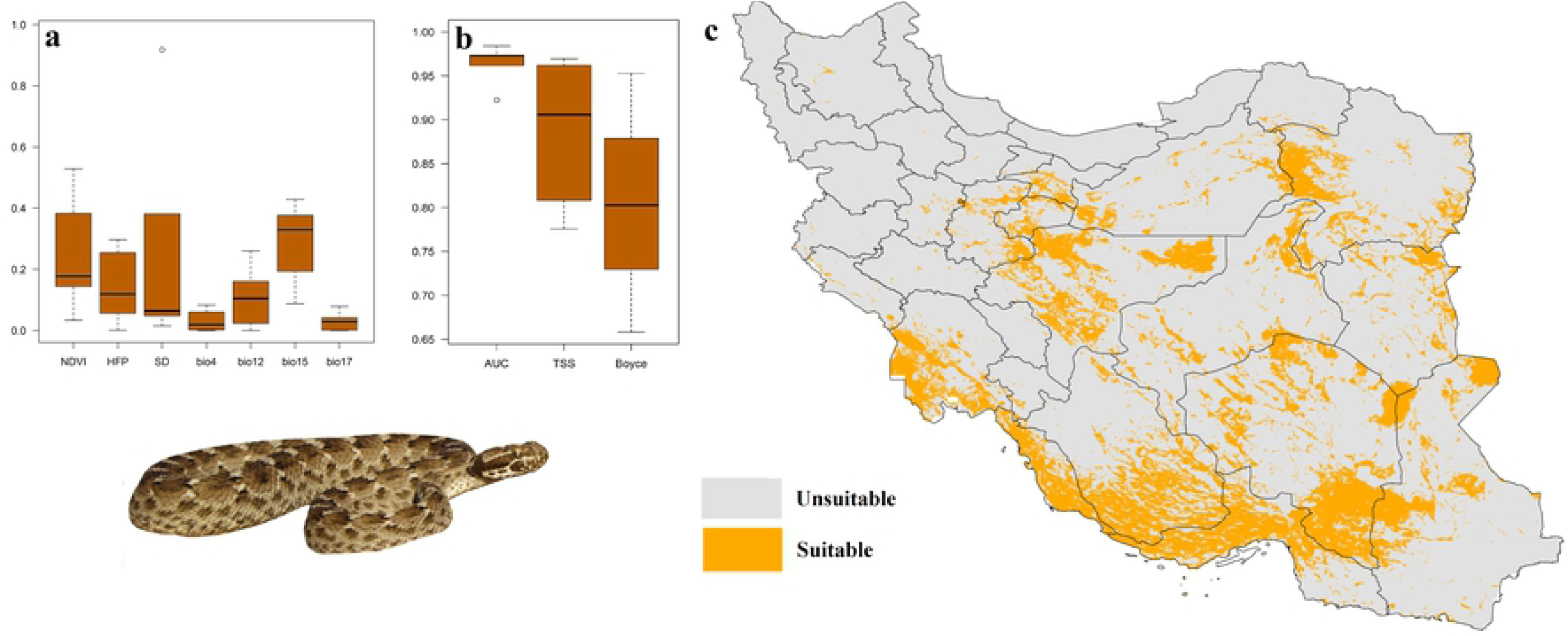
Variables importance (a), models performance (b) and habitat suitability model (c) of *Echis carinatus* in Iran.

### Habitat suitability

#### Echis carinatus

Based on ensemble model, southern part of Iran, north of Persian Gulf and vast areas in central parts of the country are identified to have highest suitability for *E. carinatus* (Fig Precipitation seasonality and NDVI and were the most important determinant of habitat suitability of the species across the Iran.

#### Macrovipera lebetina

The most suitable habitats of *M. lebetina* are located in Zagros Mountains, Alborz Mountains, Kopat-Dagh Mountains as well as in some isolated mountains in central Iran (Fig 4). Precipitation seasonality, precipitation of the driest quarter, and human footprint were the most important predictors of suitable habitats for the species.

**Fig 4.**
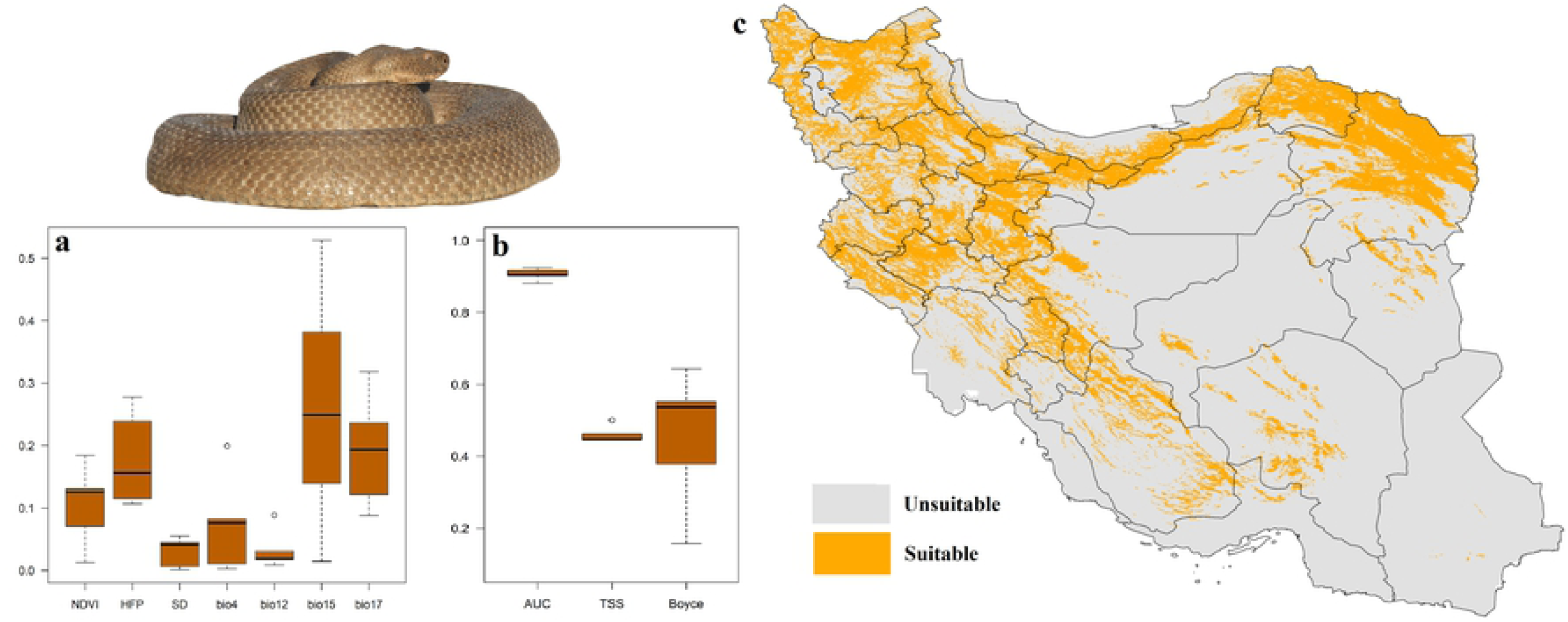
Variables importance (a), models performance (b) and habitat suitability (c) of *Macrovipera lebetina* in Iran.

#### Pseudocerastes persicus

Central, southwest and northeast of Iran have highest suitability for the *P. persicus*. While, northern parts of the country are not suitable for this species (Fig 5). Results showed that precipitation seasonality and NDVI and were the most important determinant of habitat suitability of the species across the Iran.

**Fig 5.**
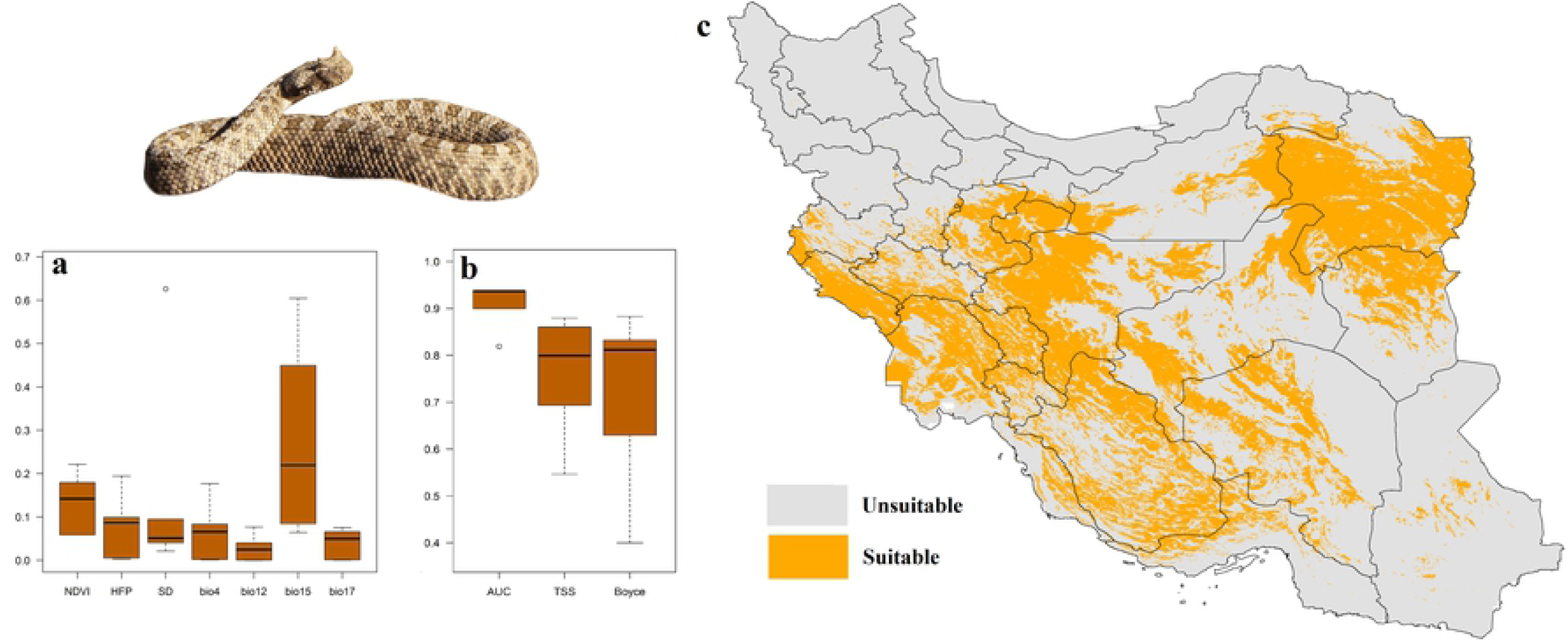
Variables importance (a), models performance (b) and habitat suitability (c) of *Pseudocerastes persicus* in Iran.

#### Naja oxiana

*Naja oxiana*’s most suitable habitats are located in north east of Iran around Kopat-Dagh Mountains as well as some isolated patches in western parts of the country (Fig 6). Precipitation of the driest quarter was the most important predictor of suitable habitats of the species.

**Fig 6.**
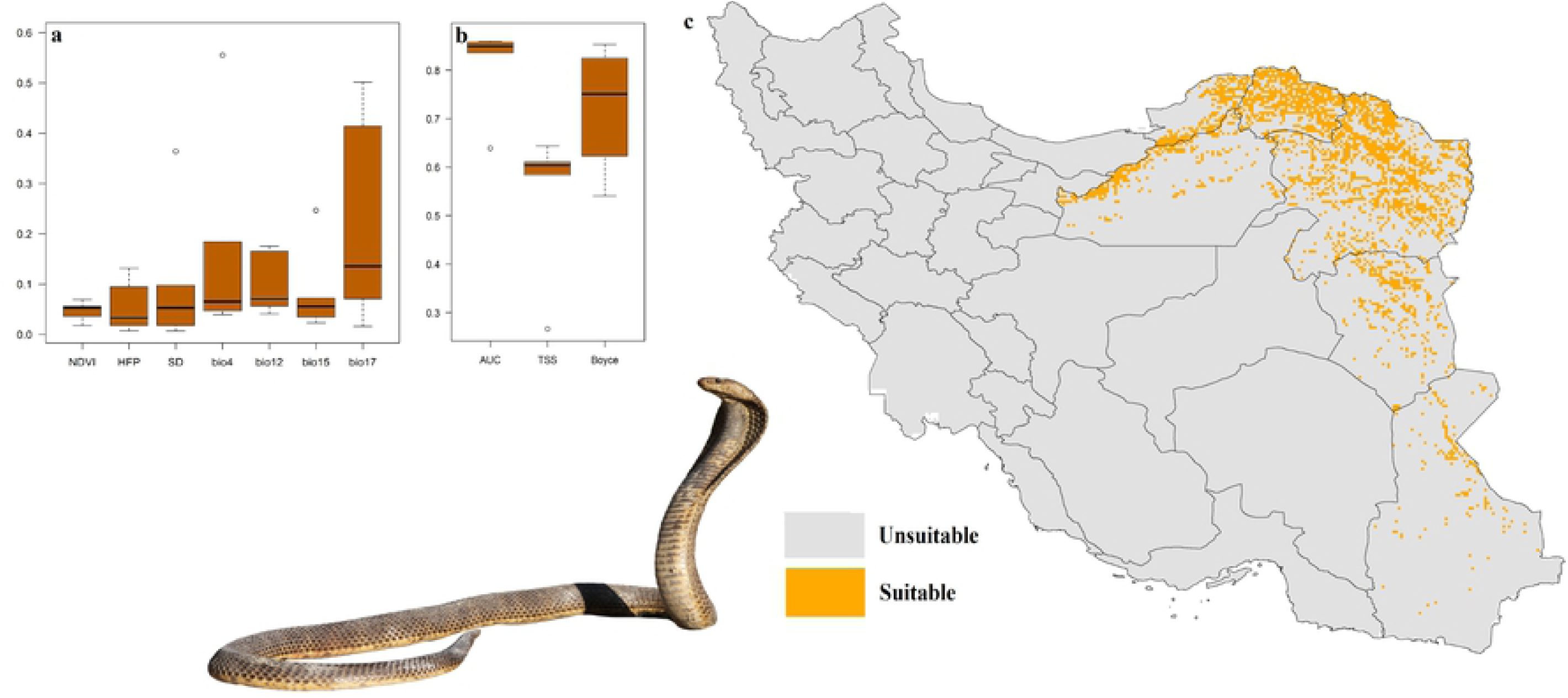
Variables importance (a), models performance (b) and habitat suitability (c) of *Naja oxiana* in Iran.

### Snakebite risk model

The four venomous snakes’ ensemble habitat suitability models were combined to develop a snakebite risk model for Iran (Fig 7). Results showed that Khorasan-e-Razavi, east of Semnan, north of Khorasan-e-Jonobi and south of Khorasan-e-Shomali provinces have highest snakebite risk in Iran. West of Khorasan-e-Razavi province has high suitability for the four venomous snakes.

**Fig 7.**
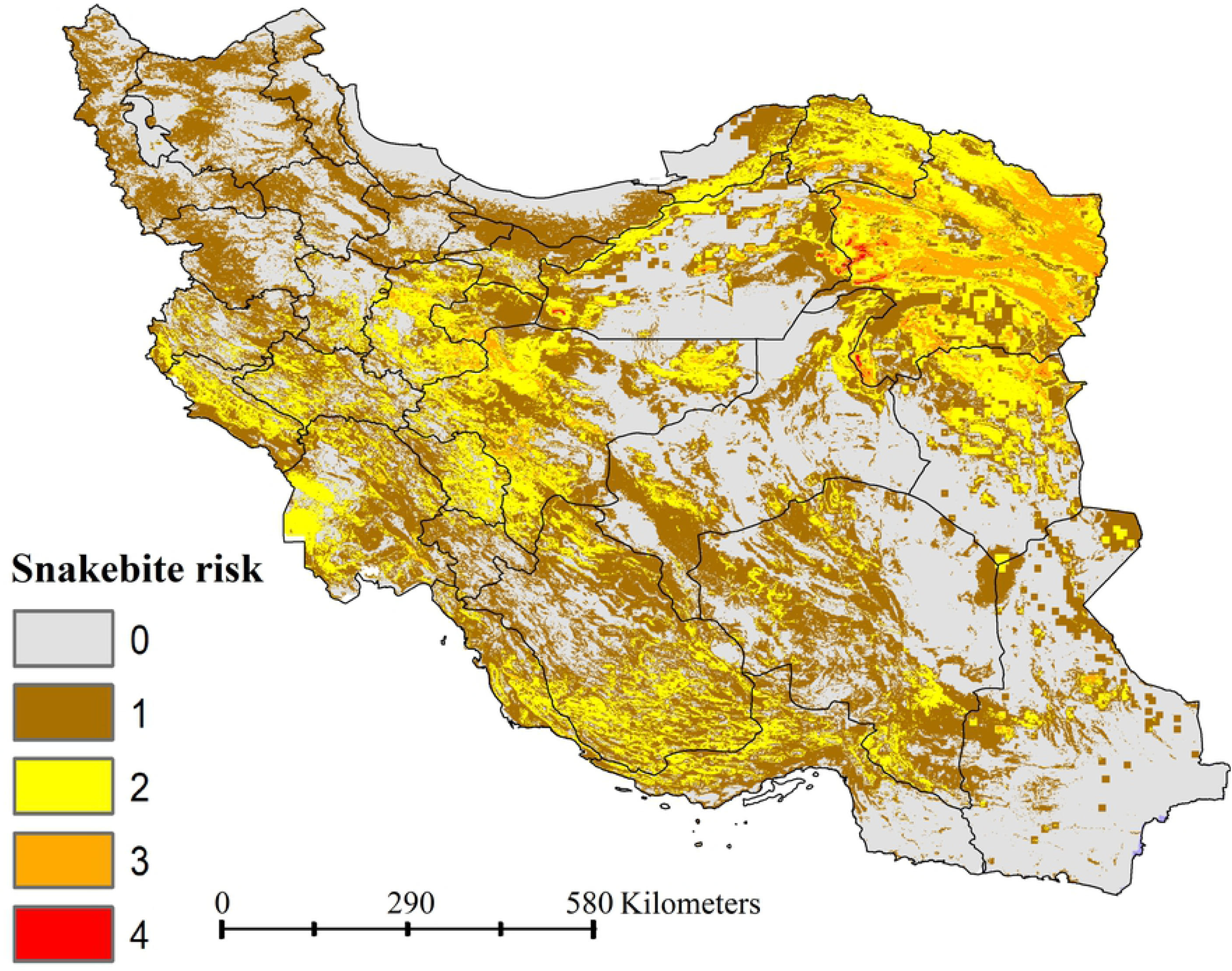
Snakebite envenoming risk model in Iran. The snakebite risk model was developed based on combined habitat suitability models’ of *Macrovipera lebetina, Echis carinatus, Pseudocerastes persicus* and *Naja oxiana*.

### Villages at risk of envenoming

Number of villages that are at the risk of envenoming by each of the four snakes (Table 2, Fig 8) were determined. Results revealed that 51,112 villages are at risk of envenoming from *M. lebetina*, 30,339 from *E. carinatus*, 51,657 from *P. persicus* and 12,124 from *N. oxiana*. Area of envenoming risk by each species was estimated (Table 2), *M. lebetina* and *N. oxiana* are identified with largest (362,558 km2) and smallest (121,803 km2) area, respectively.

**Table 2.** Area (km^2^) and number of villages that are at the snakebite risk from *Macrovipera lebetina, Echis carinatus, Pseudocerastes persicus* and *Naja oxiana* in Iran. Results are based on the ensemble models.

| Species | Villages at high risk | Villages at moderate risk | Area |
| --- | --- | --- | --- |
| *Macrovipera lebetina* | 9,824 | 41,288 | 362,558 |
| *Echis carinatus* | 5,917 | 24,422 | 321,089 |
| *Pseudocerastes persicus* | 2,150 | 49,507 | 655,415 |
| *Naja oxiana* | 3,091 | 9,033 | 121,803 |

**Fig 8.**
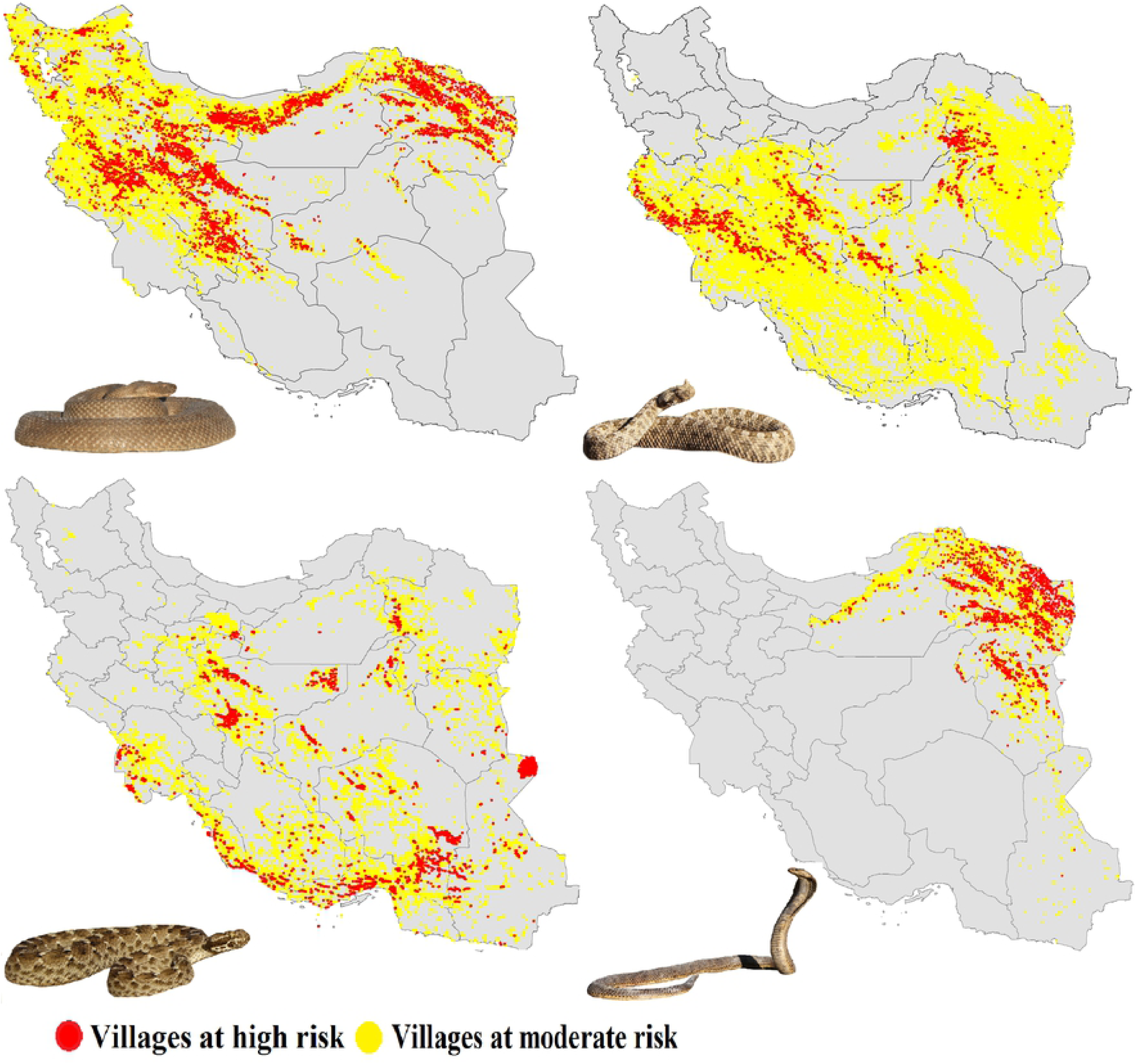
Villages at risk of envenoming from *Macrovipera lebetina, Echis carinatus, Pseudocerastes persicus* and *Naja oxiana*. Villages at high risk are shown with red circle and villages at moderate risk are shown with yellow circle.

## Discussion

With this research the first snakebite envenoming risk model was produced at fine resolution (∼1 km^2^) in Iran by modeling and multiplying habitat suitability of four medically important venomous snakes which are responsible for the most snakebite incidents in the country [18]. Northeastern parts of Iran were identified to have highest snakebite risk in the country. Results showed that thousands of villages are located in suitable ranges of the four venomous snakes, for example 51,657 villages are in the suitable range of *P. persicus*. Thus these villages and villagers are at risk of envenoming from these snakes. The snakebite risk model shows which parts of Iran are at risk of envenoming from two, three or even four of snakes. All provinces of Iran, except those in northwest of the country, are at risk of envenoming from at least two venomous snakes. This highlights importance of snakebite envenoming as public health problem in Iran.

Under climate change some venomous species may expand their distribution ranges [56-86], thus, envenoming risk will likely vary [12, 17, 87]. For instance, Nori et al. [12] modeled distribution of five venomous snakes in Argentina for 2030 and 2080. They found that the snakes’ suitable climate spaces will increase in human populated areas of the country. In another study, Zacarias & Loyola [57] modeled current and future distributions of 13 snakes in Mozambique and showed that venomous snake distribution will change under climate change. They concluded patterns of snakebite risk may change due to climatic changes [57]. Results of current research revealed that climatic variables play an important role in shaping the distribution of four venomous snakes in Iran, thus their distribution may alter with changing climate. It is predicted that suitable habitats of *Echis carinatus* will increase in Iran [88]. This species is an important source of snakebite in the country and its distributional range will likely increase under climate change putting more populations and human settlements at risk of envenoming by this snake until 2070.

Venomous snake populations are declining and many of them are listed by the IUCN Red List as Vulnerable, Endangered or Critically Endangered [89]. Conservation of snakes especially venomous snakes is a big challenge [89] as it is not easy to convince people to conserve venomous snakes which are a significant cause of human mortality and morbidity [4, 6, 14, 15]. It is necessary to identify areas with high risk of snakebite envenoming and prioritize those areas for snakebite risk management in each country. In this study potential risk areas were identified by using habitat suitability models as an indicator of snakebite risk. Future snakebite monitoring in areas with high risk can show the model performance and its power in predicting envenoming events across the country.

Results of this study can be used to reduce snakebite and venomous snake conflicts with local people, farmers and shepherds in the country. Snakebite risk can be reduced through community education [90-92]. We encourage education of local people about snakebite prevention measures in areas with highest snakebite risk. There are simple solutions to prevent snakebite envenoming in areas with high risk, like villages in west of Khorasan-e-Razavi province. For example, using bed nets and protecting feet, ankles and lower legs by wearing boots can significantly reduce snakebite envenoming [91-93]. We also suggest that areas where snakebite envenoming risk is high should be monitored to determine envenoming events and villagers in these areas must always have access to antivenom supplies [14].

Species distribution models are becoming important tools in public health research [12, 94-103]. We encourage public hearth researchers to apply species distribution models in developing snakebite risk map using venomous snakes’ habitat suitability as an indicator, especially in data poor regions of the world [27, 52-54]. Our approach has the potential for practical application in other countries whit high snakebite risk.

## Acknowledgments

We thank Ollie Thomas for improving the English of the manuscript.

